# Effects of carbon dioxide enrichment and environmental factors on photosynthesis, growth and yield and their interaction in cucumber: a meta-analysis

**DOI:** 10.1101/2025.10.31.685732

**Authors:** Xin Liu, Xinying Liu, Yaliang Xu, Zheng Wang, Qiying Sun, Sujun Liu, Binbin Liu, Qingming Li

## Abstract

Despite the ongoing increase in atmospheric carbon dioxide concentrations, these levels remain well below the optimal threshold for cucumber growth and development. In addressing the challenges posed by climate change to agricultural production, the use of carbon dioxide enrichment (eCO_2_) has become increasingly widespread in order to meet global demand for cucumbers. Nevertheless, the impact of eCO_2_ and environmental factors on cucumber growth remains to be elucidated. In this study, we conducted a meta-analysis of 73 research papers on eCO_2_ in cucumbers was conducted, and their photosynthesis, growth, and yield were analysed. In summary, under conditions of elevated CO_2_ levels, the net photosynthetic rate, biomass, and yield of cucumbers exhibited a marked increase of 56.31%, 27.75%, and 21.98%, respectively. Concurrently, stomatal conductance and transpiration rate exhibited a decline of 36.07% and 30.42%, respectively. In the context of the implementation of eCO_2_ at varying levels within a production environment, the prevailing recommendation pertains to the range of 800-1200 ppm. It is advised that this be integrated with elevated light intensity, augmented temperature, suitable humidity levels, and a sufficient supply of fertiliser to achieve a synergistic effect. The findings of this study contribute to enhancement of environmental control in the context of greenhouse cultivation of cucumber in response to climate change, as well as promoting the green and sustainable development of the cucumber industry.

## 1. Introduction

The global climate system is undergoing unprecedented and profound changes, and ongoing climate change has become a core challenge constraining sustainable agricultural production. According to the Sixth Assessment Report released by the Intergovernmental Panel on Climate Change (IPCC)(IPCC, 2023) and the research findings of Horton et al.(Horton et al., 2021), the sharp rise in atmospheric carbon dioxide concentrations is one of the most significant features of current climate change. CO□ serves as the core raw material for plant photosynthesis, playing an irreplaceable role in carbohydrate synthesis, energy conversion, and material metabolism. Despite atmospheric carbon dioxide concentrations having historically exceeded 400 ppm, research indicates that this concentration level is far from optimal for the growth and development of most vegetable crops(Syed and Hachem, 2019). In scenarios involving controlled environment agriculture, shortages of carbon dioxide are particularly evident. Due to the enclosed nature of facility environments and limitations in air circulation, the carbon dioxide circulation system within facilities differs significantly from that of the external atmosphere. This sustained low-concentration environment leaves facility vegetables in a state of carbon starvation, severely limiting the accumulation of photosynthetic products and plant growth and development, thereby becoming a key bottleneck constraining the quality and efficiency of facility agriculture(Zhang *et al*., 2014). To address the challenges posed by climate change to agricultural production whilst meeting the growing global demand for vegetable supply, carbon dioxide fertilisation technology has been gradually applied in horticultural facilities since the early 20th century. Following a period of over a century characterised by technological iteration and practical exploration, this technology has evolved into a significant means of enhancing the yield and quality of greenhouse vegetables(Dong *et al*., 2018; Hao *et al*., 2020).

In the global vegetable industry landscape, cucumbers have always been of significance due to their diverse consumption scenarios and massive market demand. The global area under cucumber cultivation has been steadily increasing, making it one of the core drivers for ensuring stable vegetable supply and promoting agricultural economic development(Zhao *et al*., 2019). However, in the current context of increasingly widespread protected cultivation, cucumber production faces numerous challenges posed by environmental constraints. While controlled environment agriculture provides relatively controllable conditions such as temperature, humidity, and light for cucumber growth, the enclosed nature of such spaces leads to imbalances in carbon dioxide circulation. As a typical C3 plant, the Calvin cycle in cucumber leaves relies on carbon dioxide as a carbon source and is highly sensitive to changes in carbon dioxide concentration(Chen *et al*., 2022). In facility cultivation scenarios, after sunrise, as photosynthesis in plants rapidly begins, carbon dioxide concentrations inside the facility drop sharply within a short period of time(Zhang *et al*., 2020). At this point, the carbon dioxide concentration within the facility environment is far below the atmospheric background value, causing the net photosynthetic rate of cucumber leaves to decrease by approximately 35-40%(Chen *et al*., 2019). Prolonged exposure to this low-carbon dioxide environment has been demonstrated to significantly reduce the carbon assimilation capacity of cucumber plants, directly leading to a decrease in cucumber yield of approximately 20-30%, while the nutrient content of the fruit is also negatively impacted to varying degrees(Kläring *et al*., 2007).

It is generally accepted that eCO_2_ have a positive impact on photosynthetic rate and yield(Li *et al*., 2025). Carbon dioxide enrichment of cucumbers at concentrations of 400-1600 ppm can increase photosynthetic rate and yield of cucumbers to varying degrees(Liu *et al*., 2018; Abdeldaym *et al*., 2024). The increase in yield was attributed to the increase in net photosynthetic rate and water utilisation, which promoted leaf growth, increased plant yield, and plant tolerance to stressful environments such as heat and drought(Huang and Xu, 2015). However, the effects of eCO_2_ do not exist in isolation, and there are complex interactions between its effects and environmental factors such as light intensity, humidity, and temperature. Other environmental factors such as light and humility affect the effect of eCO_2_ on cucumber, while, at the same time, eCO_2_ interacts with other environmental effects on cucumber(Long *et al*., 2006). Therefore, the synergistic and antagonistic relationships between eCO_2_ and environmental factors make the response of cucumber to CO□A1enrichment highly complex and dynamic.

A significant number of studies have focused on food crops such as wheat and soybeans(Ziska and Bunce, 2007; Wang *et al*., 2013). However, limited number of studies have conducted comprehensive analyses of the effects of eCO_2_ on vegetables, and there is an even greater lack of specific analyses on cucumbers alone. Despite the plethora of reports on the effects of eCO_2_ on cucumber growth, systematic studies on the coupled effects of different eCO_2_ concentrations with environmental factors remain insufficient. In particular, the regulatory mechanisms of cucumber physiological metabolism under the interactive effects of eCO_2_ with factors such as light, water, and temperature remain unclear. Consequently, a thorough literature review is imperative to elucidate the effects of eCO_2_ on cucumber growth, photosynthesis, and yield, as well as its interactive effects with the environment. This not only contributes to refining the theoretical framework for environmental regulation in protected cucumber cultivation but also provides a theoretical basis and practical guidance for the development of precise and intelligent CO□ fertilisation techniques. This holds significant scientific and practical value for promoting the green and sustainable development of the cucumber industry.

## 2. Materials and methods

### 2.1 Data collection

We searched for peer-reviewed journal articles using the “China National Knowledge Infrastructure”, “Web of Science”, “Science Direct”, and “Google Scholar” (as of January 2025), with the following keyword combinations were used: (elevated carbon dioxide OR eCO_2_ OR CO_2_ enrichment OR carbon dioxide enrichment) and (cucumber). Articles must be selected as eligible studies based on the following criteria:(i) Research paper. (ii) The study included both control and CO_2_ enrichment treatments. (iii) At least one variable was recorded including plant height, Stem diameter, leaf area, biomass, photosynthesis indicators and yield. (iv) Each treatment was replicated at least three times. After a second screening by reading the titles and abstracts of the literature and excluding those that did not meet the selection criteria, the retained literature was subjected to a more rigorous screening. The entire flowchart of the review programme is shown in Fig. S1. Publications and datasets can be found in Table S1. Finally, 73 articles from around the world met our criteria and were used in our meta-analysis (Fig. 1).

**Fig. 1.**
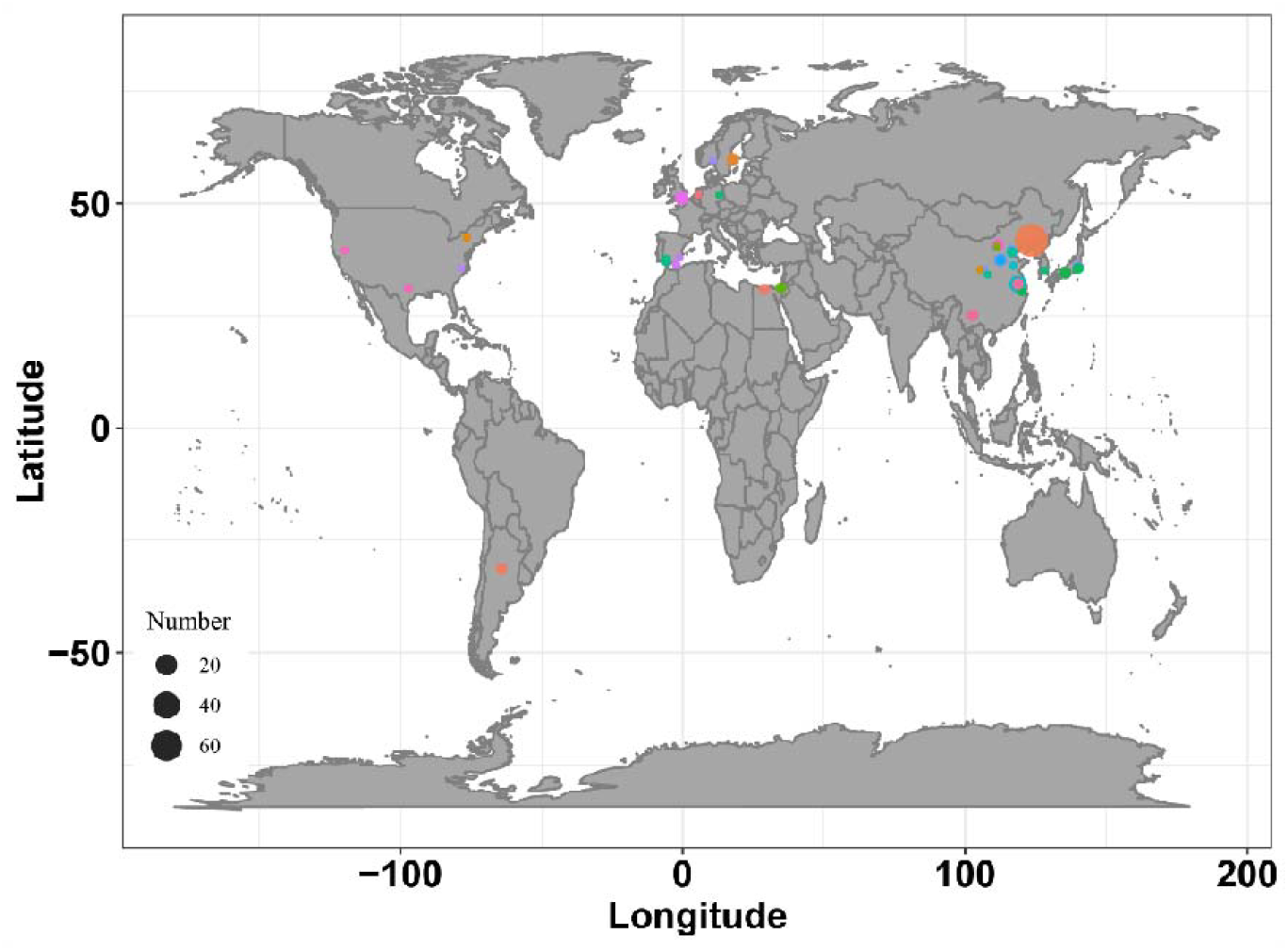
Spatial distribution of published papers, with locations represented by points. The size of the points represents the size of the sample size for different studies.

We further extracted light intensity, temperature, humidity, and concentrations of nitrogen (N), phosphorus (P) and potassium (K) fertilisers in the nutrient solution to assess the effects of CO_2_ enrichment on cucumber growth, photosynthesis, and yield. All target variables (mean, standard error, sample size) were displayed graphically or numerically and data presented graphically were extracted using WebPlotDigitizer(Burda *et al*., 2017). In instances where the paper did not mention the standard deviation (SD) or standard error (SE) of the variable, the metagear package was utilised to estimate the SD(Lajeunesse, 2016)).

### 2.2 Meta-analysis

Effect sizes were calculated using the natural logarithm of response ratios (lnRR) according to the meta-analysis method(Hedges *et al*., 1999). The calculation formula was as follows (1):

*X*_*t*_ and *X*_*c*_ are the mean values of treatment and control, respectively. The variance (V_lnRR_) for each effect size was calculated as follows (2):

Where *n*_*t*_ and *n*_*c*_ are the sample sizes of the treatment and control groups, respectively, and *s*_*t*_ and *s*_*c*_ are the standard deviations of the treatment and control groups, respectively.

Weighted averages are used to produce maximum accuracy and can eliminate as much variation as possible. The weighted average response ratio (lnRR_+_) and weights (W_i_) were calculated from (3) and (4), respectively.

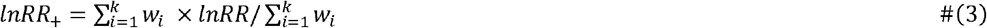

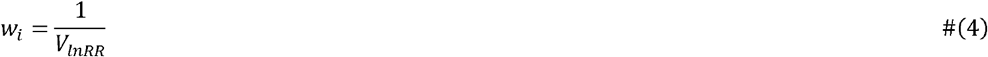

Here *i* and *k* denote the number of comparisons and cumulative studies, and *w*_*i*_ denotes the weight of independent studies. Weighted effect ratio variance (Var(lnRR_+_)) and 95% confidence intervals (95% CI) were calculated using equations (5), (6).

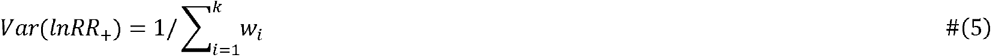

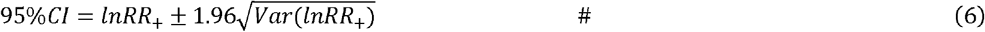

For ease of interpretation of the analysis, weighted effect sizes were converted to percentage changes (%) using equation (7) based on comparisons between the experimental and control groups.

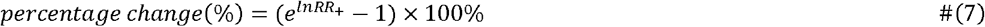

### 2.3 Data-analysis

Meta-analyses, statistical analyses and plots were performed in R (version 4.4.1) software. Response ratios were calculated using a random-effects model with restricted maximum likelihood (RMEL) to calculate weighted response ratios and heterogeneity between the data pairs (I^2^)(Wang *et al*., 2024). We used Random Forest to examine the relative importance of each predictor variable to the response ratio. The potential for publication bias to influence the outcomes of meta-analyses necessitates the implementation of fail-safe analyses to identify any such bias(Du *et al*., 2023). The results demonstrate that Rosenthal > 5N + 10 (N representing the sample number) and I^2^ are greater than 95% (Table S1). This indicates that this study was not affected by publication bias and that the conclusions are reliable.

## 3. Results

### 3.1 Effect of eCO_2_ on photosynthesis and water use efficiency in cucumber

The findings of the present study demonstrated that the application of elevated CO□ levels resulted in a significant augmentation in the net photosynthetic rate and water use efficiency of cucumber, with respective increases of 56.31% (95% CI: 50.31% to 62.32%) and 121.11% (95% CI: 86.26% to 155.96%). Concurrently, stomatal conductance and transpiration rate exhibited a decline, with reductions observed of 36.07% (95% CI: −47.31% to −24.83%) and 30.42% (95% CI: −39.20% to −21.64%), respectively (Fig. 2). The eCO_2_ environment was found have a stimulatory effect on the net photosynthetic rate, with this effect becoming more pronounced as the concentration increased (Table S2). It was demonstrated that the application of eCO_2_ led to a reduction in the transpiration rate. However, the differences observed between the eCO_2_ gradients were not found to be statistically significant. Notwithstanding, these results led to a significant increase in water use efficiency, by almost 150% at 400-800 ppm and 1200-1600 ppm.

**Fig. 2.**
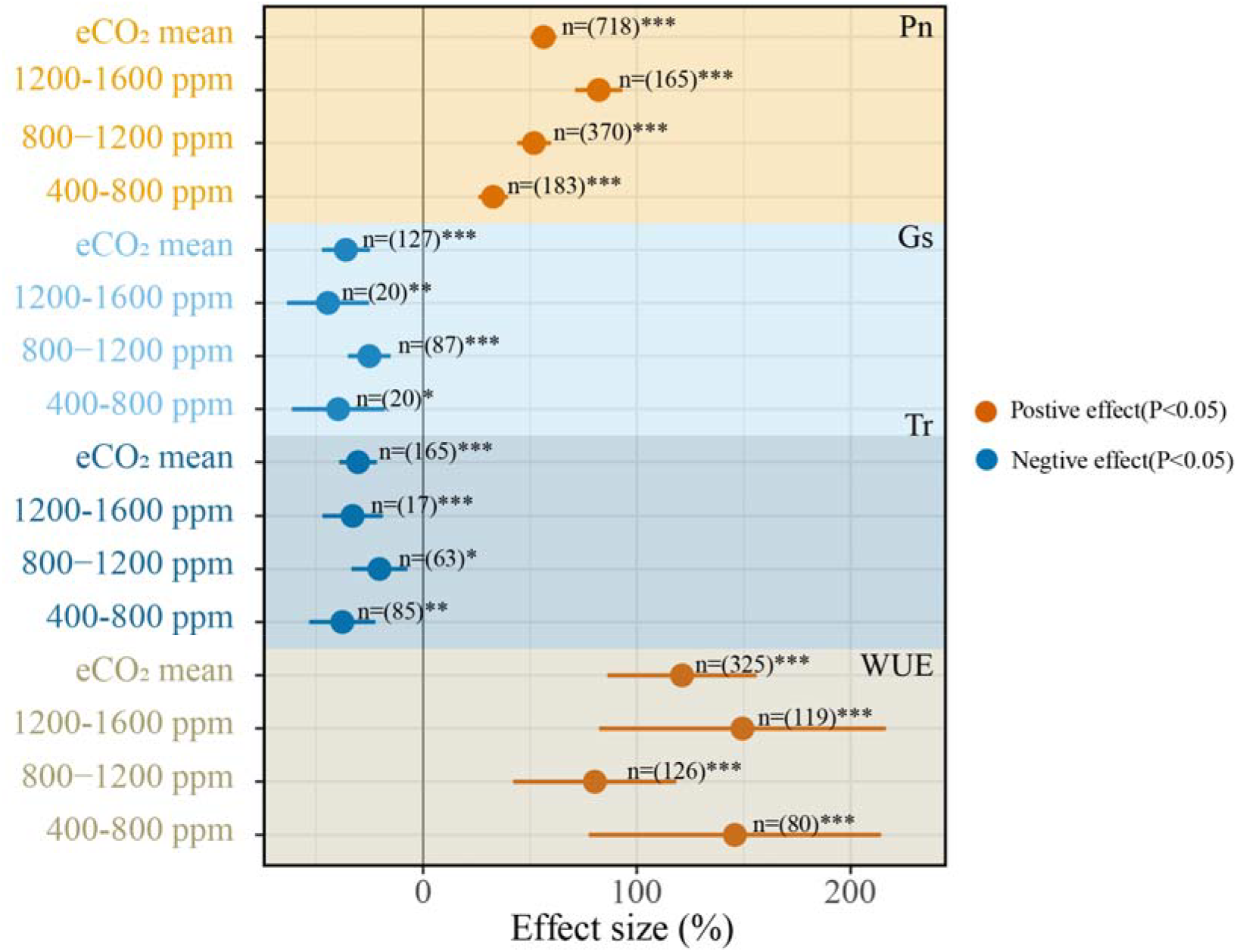
Overall response rate of cucumber photosynthesis and water use efficiency to eCO_2_. The points represent the mean values and the numbers next to them indicate the cumulative sample number of studies. Error lines represent 95% CIs. Asterisks (*) indicate significant differences (*: p < 0.05, **: p < 0.01, ***: p < 0.001). The same as above.

### 3.2 Interaction between eCO_2_ and environmental factors on cucumber photosynthesis

The net photosynthetic rate of cucumber exhibited an upward trend in conjunction with the CO_2_ concentration (Fig. S2). An analysis of the environmental factors under eCO_2_ was conducted, revealing that elevated light intensity, temperature, humidity and N and K supply were more effective in increasing the net photosynthetic rate, particularly at 1200-1600 ppm. (Fig. 3). As demonstrated in Figure S3, the net photosynthetic rate was found to be significantly influenced by temperature, light intensity, and nitrogen supply. A subsequent analysis of these factors revealed an increasing trend in temperature and light intensity, followed by a subsequent decreasing trend. In contrast, an increase in N supply was found to result in a consistent increase in the net photosynthetic rate (Fig. S4).

**Fig. 3.**
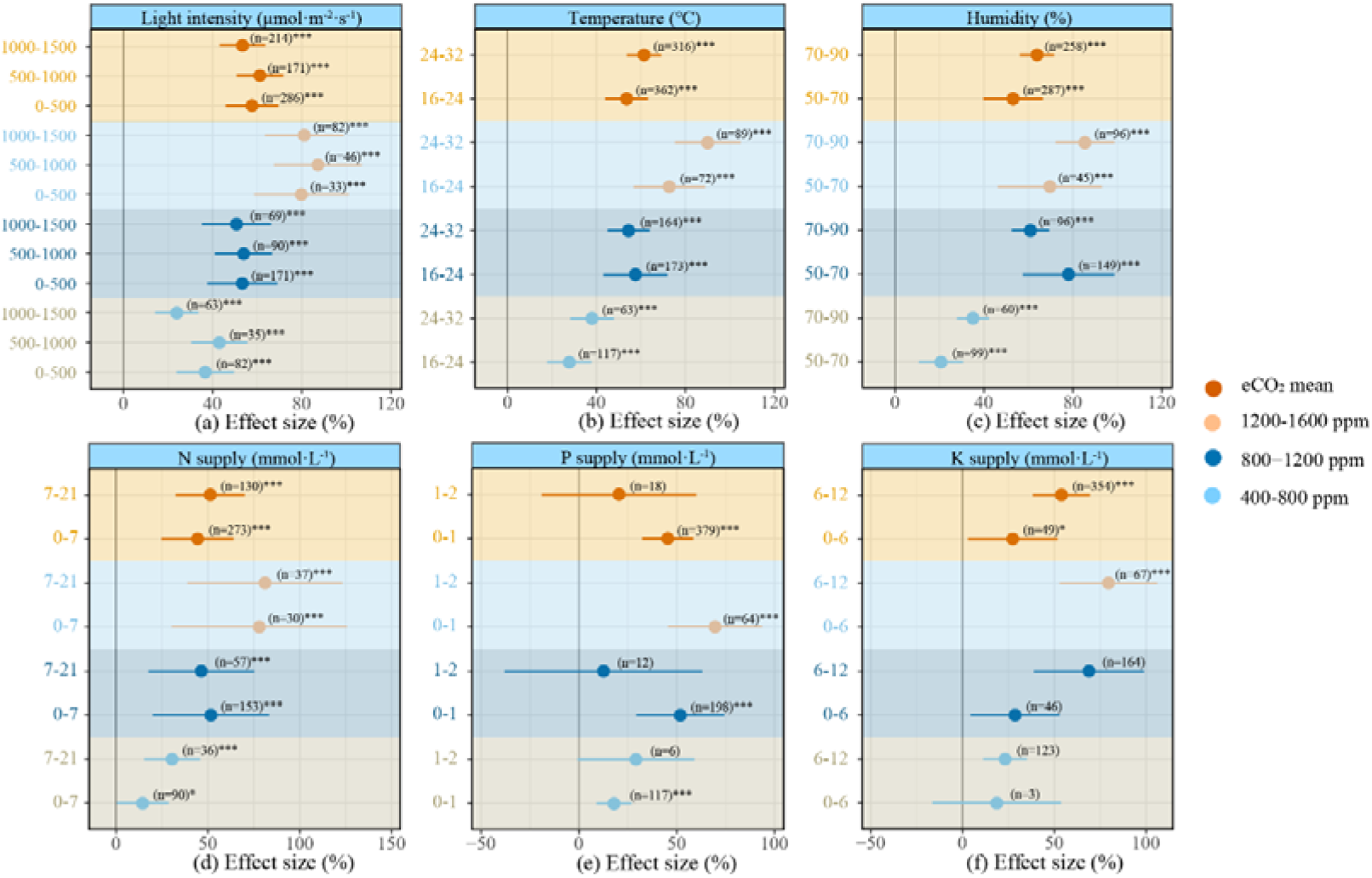
Effect of various environmental factors on cucumber biomass accumulation. The points represent the mean values and the numbers next to them indicate the cumulative sample number of studies. Error lines represent 95% CIs. Asterisks (*) indicate significant differences (*: p < 0.05, **: p < 0.01, ***: p < 0.001). The same as above.

### 3.3 Effect of eCO_2_ on cucumber biomass accumulation and allocation

As demonstrated in Figure 4, eCO_2_ resulted in a substantial increase in cucumber biomass accumulation. demonstrated a clear increase in the biomass of cucumber roots, stems and leaves, with eCO_2_ increasing the biomass by 26.92% (95% CI: 17.05% to 36.80%), 17.79% (95% CI: 5.70% to 29.88%), 22.48% (95% CI: 10.55% to 34.42%), respectively. The mean CO□ concentration of 930 ppm was found to have a significant positive effect on the total biomass accumulation of cucumber, with an increase of 27.75% (95% CI: 20.89% to 34.61%). In eCO_2_ treatments, 800-1200 ppm ware able to significantly increase the aboveground and total biomass accumulation of cucumber. While higher CO_2_ concentration favoured the accumulation of belowground biomass. It is worthy of note that, at 400-800 ppm however, reduced stem biomass accumulation (Table S2).

**Fig. 4.**
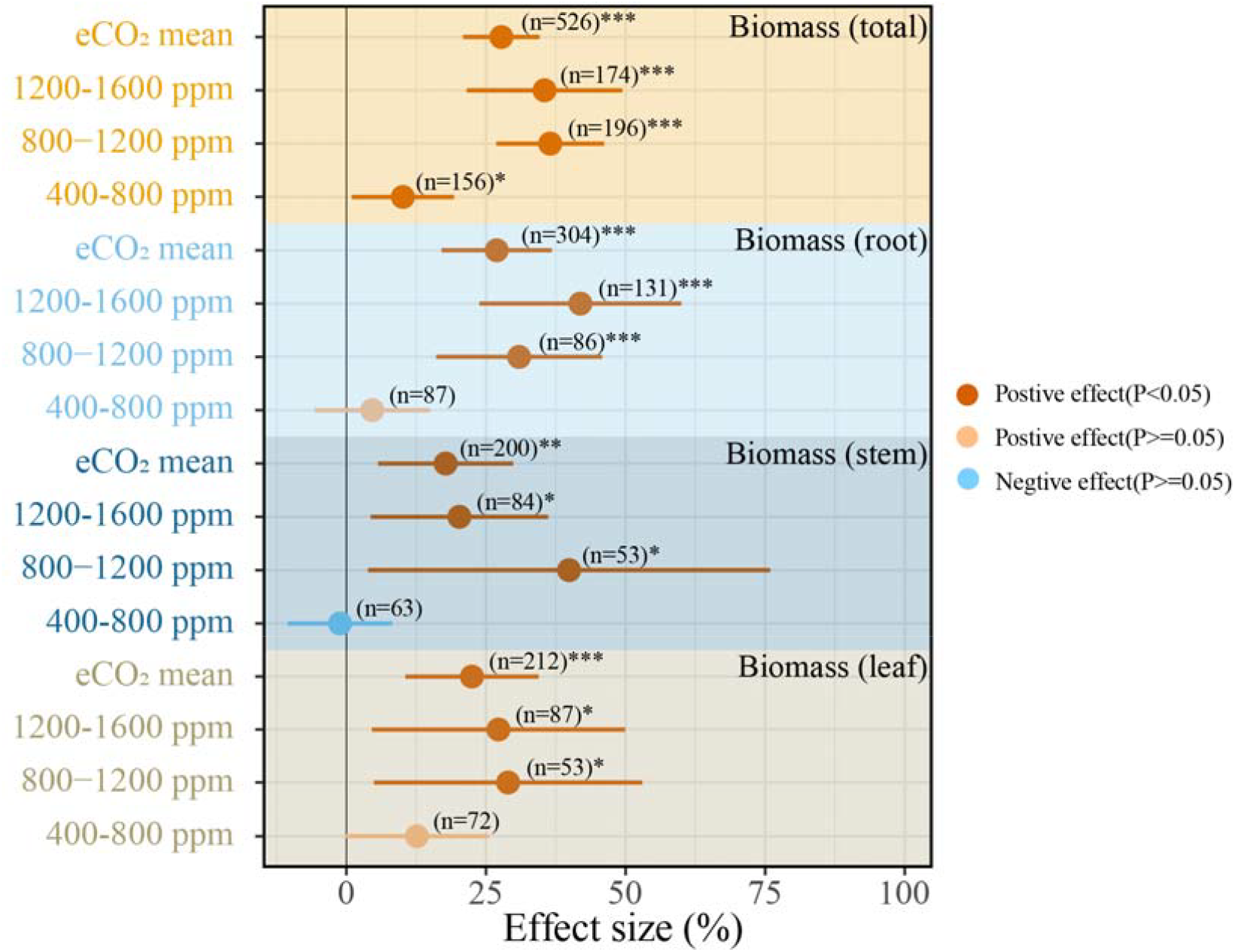
Overall response rate of cucumber biomass accumulation to eCO_2_. The points represent the mean values and the numbers next to them indicate the cumulative sample number of studies. Error lines represent 95% CIs. Asterisks (*) indicate significant differences (*: p < 0.05, **: p < 0.01, ***: p < 0.001). The same as above.

### 3.4 Interaction between eCO_2_ and environmental factors on cucumber biomass accumulation

As the concentration of CO□ increased, there was a concomitant increase in cucumber biomass accumulation (Fig. S5). An analysis of the interaction of eCO_2_ and environmental variables found that higher light intensity interacted with different eCO_2_ all showed stronger effects on increasing biomass; while humidity showed the opposite result (Fig. 5). By weighting analysis, temperature, N supply, and humidity were important regulators affecting biomass accumulation (Fig. S6). Under eCO_2_, a gradual increase in biomass was accompanied by an increase in light intensity, temperature, and N supply, while humidity, P, and K supply showed a decreasing trend (Fig. S7).

**Fig. 5.**
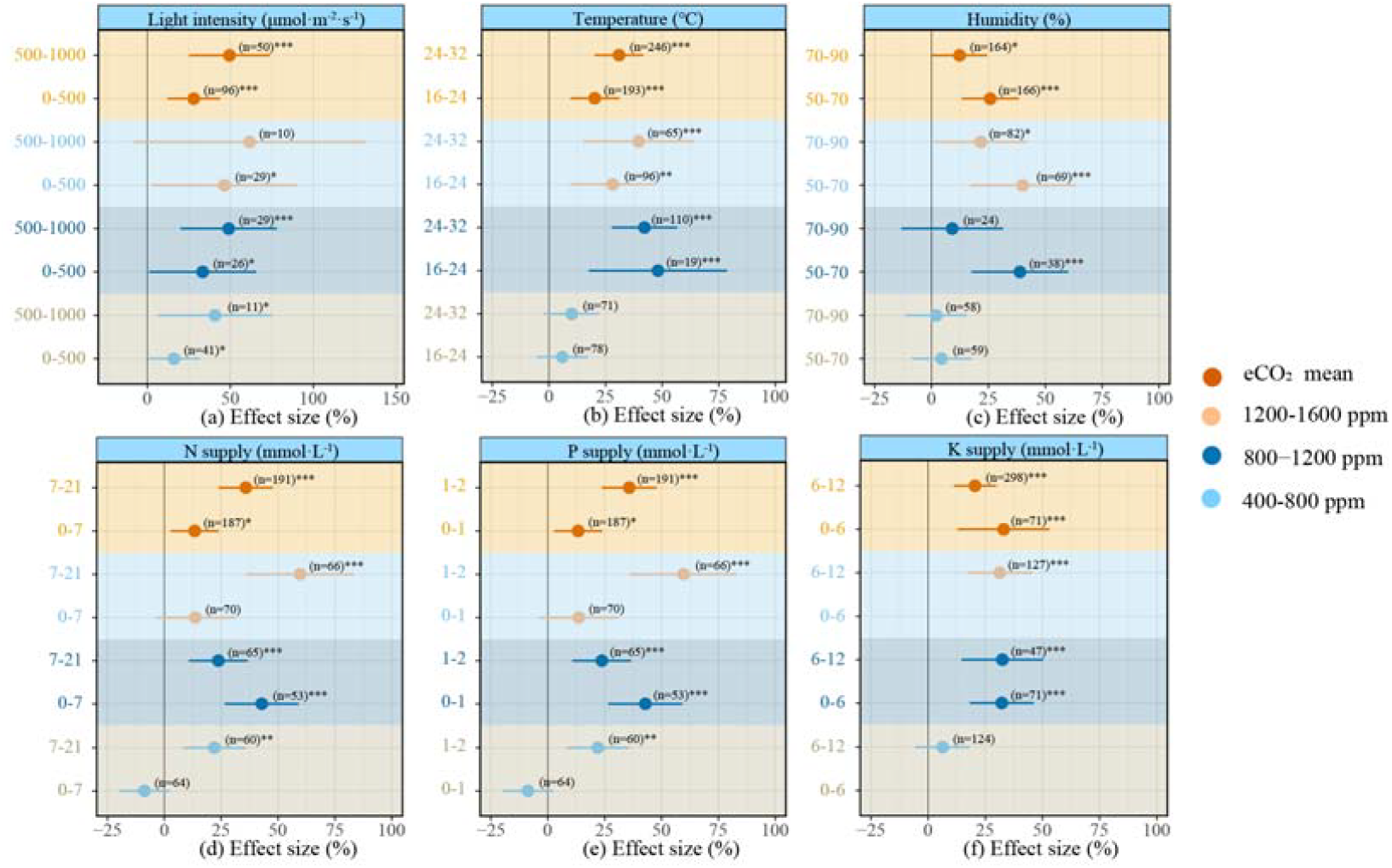
Effect of various environmental factors on cucumber biomass accumulation. The points represent the mean values and the numbers next to them indicate the cumulative sample number of studies. Error lines represent 95% CIs. Asterisks (*) indicate significant differences (*: p < 0.05, **: p < 0.01, ***: p < 0.001). The same as above.

### 3.5 Effect of eCO_2_ on cucumber growth and yield

The consolidation of all the eCO_2_ data yielded results that indicated a significant impact of eCO_2_ on the growth and yield of cucumber (Fig. 6). The results demonstrated a significant increase in plant height, stem diameter and leaf area of cucumber under eCO_2_, with increases of 19.39% (95% CI: 10.62% to 28.16%), 11.46% (95% CI: 7.41% to 15.51%) and 27.19% (17.76% to 36.61%), respectively. As the concentration of CO□ increased under eCO_2_ conditions, there was a significant increase in the height of the plants, the diameter of the stems, and the area of the leaves of the cucumbers (Fig. S8, Table S2). Cucumber yield was highest at 800-1200 ppm, with an average increase in CO_2_ to 970 ppm, resulting in a 21.98% (95% CI: 16.60% to 27.36%) increase in yield.

**Fig. 6.**
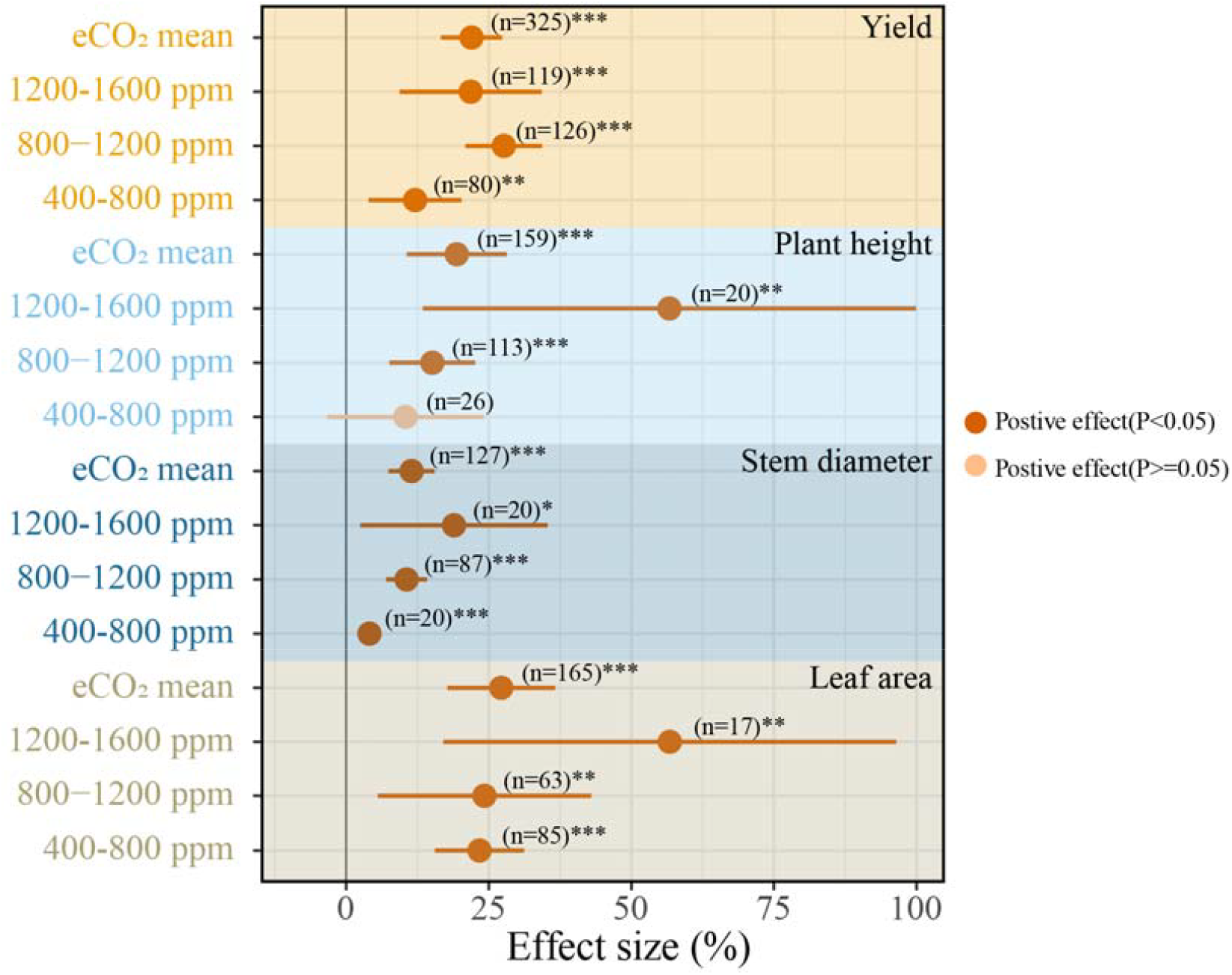
Overall response rates of cucumber plant height, stem diameter, leaf area and yield to eCO_2_. The points represent the mean values and the numbers next to them indicate the cumulative sample number of studies. Error lines represent 95% CIs. Asterisks (*) indicate significant differences (*: p < 0.05, **: p < 0.01, ***: p < 0.001).

### 3.6 Interaction between eCO_2_ and environmental factors on cucumber growth and yield

Other environmental factors also affect vegetable yields to varying degrees when vegetables are grown in eCO_2_ environments(Dong *et al*., 2020). The findings of the present meta-analysis confirmed that the effect of eCO_2_ on cucumber yield varies to some extent with different levels of various factors, including light intensity, temperature, humidity and the supply of N, P and K (Fig. 7). According to the Random Forest analysis of the weights of environmental factors interacting under eCO_2_, temperature, humidity, and light intensity are important regulators of cucumber yield (Fig. S9). Higher light intensity, temperature, and N supply were more effective in promoting cucumber yield, whereas excessive humidity, P, and K supply were not (Fig. S10). It is evident that an optimal level of CO□ concentration, ranging from 800 to 1200 ppm, is conducive to enhancing yield in the context of environmental and eCO_2_ interactions.

**Fig. 7.**
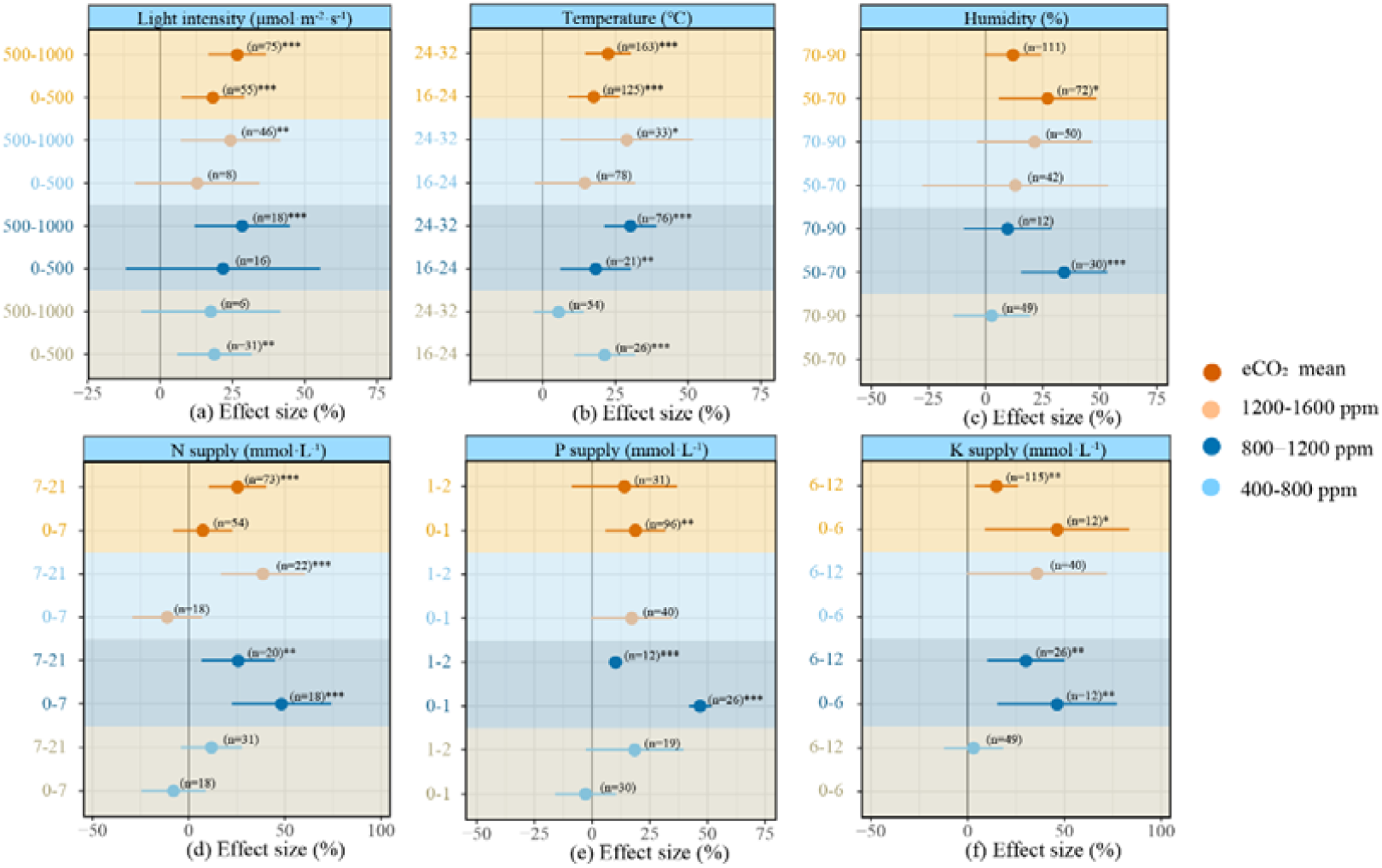
Effect of various environmental factors on cucumber yield under eCO_2_. The points represent the mean values and the numbers next to them indicate the cumulative sample number of studies. Error lines represent 95% CIs. Asterisks (*) indicate significant differences (*: p < 0.05, **: p < 0.01, ***: p < 0.001). The same as above.

## 4. Discussion

### 4.1 The effect of carbon dioxide concentration on cucumber photosynthesis, growth and yield

Increases in atmospheric carbon dioxide concentrations have the potential to positively impact agricultural production, particularly in the future. A comprehensive understanding of the growth response of cucumbers to eCO_2_ is imperative for sustainable development. The findings of this study, as demonstrated by the meta-analysis, indicated a robust and positive correlation between elevated carbon dioxide concentrations and the plant height, stem thickness, and leaf area of cucumbers (Fig. S8). It is evident that eCO_2_ exerts a favourable influence on the growth and yield of cucumbers. It has been demonstrated that an increase in carbon dioxide concentrations results in elevated levels of carbon dioxide in plant leaves, thus enhancing photosynthetic efficiency. Plants are able to convert carbon dioxide into organic compounds such as sugars more quickly, providing more energy and nutrients for plant growth(Sugiura *et al*., 2024). At the same time, increased carbon dioxide concentrations typically lead to a decrease in stomatal conductance, reducing transpiration and water loss, and thereby improving water use efficiency(Robredo *et al*., 2007; Osman *et al*., 2024). The enhancement in photosynthesis and water use efficiency observed in cucumbers cultivated under elevated CO_2_ conditions is the primary factor contributing to the substantial increases in plant height, stem diameter, leaf area, and yield (Fig. 8).

**Fig. 8.**
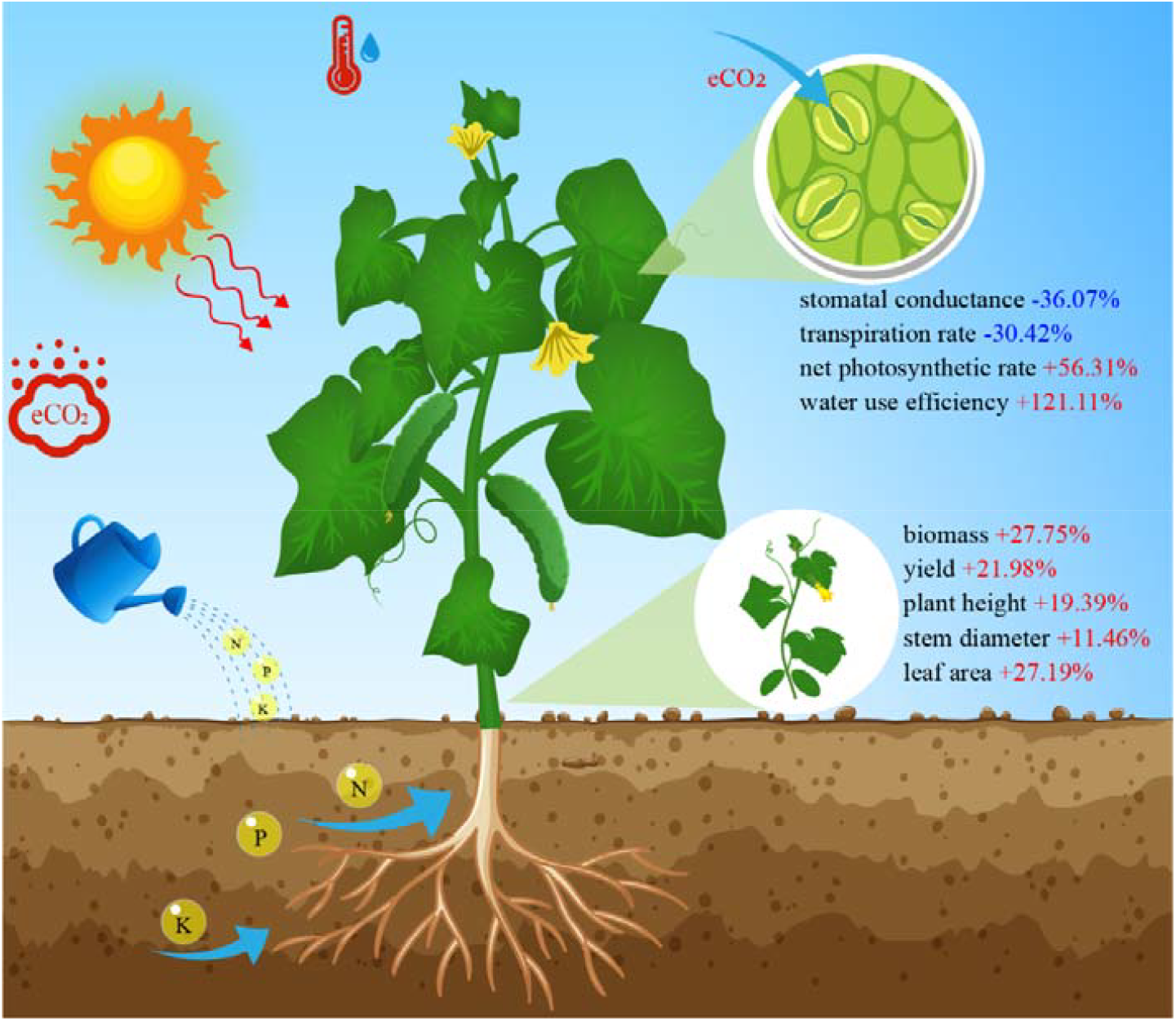
Schematic representation outlining changes in cucumber growth, photosynthesis and yield under eCO_2_.

Research indicates that eCO_2_ can enhance photosynthetic efficiency, enabling plants to fix more carbon and thereby increase biomass(Qiang *et al*., 2020; Hou *et al*., 2021). As the levels of carbon dioxide in the atmosphere increase, there is a concomitant and significant increase in the overall biomass accumulation of cucumbers. However, at carbon dioxide concentrations of 400-800 ppm, this effect appears to be less favourable for stem biomass accumulation (Fig. 4). This finding suggests that the promotional effect may be subject to variation and may be influenced by various factors, including plant species, growth conditions, and the specific experimental design(Maschler *et al*., 2022). Whilst acknowledging the constant interplay between eCO_2_ and other environmental factors, it is vital to recognise that these interactions do not inherently negate the overall promotional effect of eCO_2_ on cucumbers. It is important to note that slight variations in the effects observed under different environmental conditions may be attributable to these interacting factors.

### 4.2 The effect of light intensity on cucumber photosynthesis, growth and yield

The effects of CO_2_ enrichment on plants are not independent of other environmental factors, but rather interact with them. In the context of CO□ enrichment, it has been demonstrated that the net photosynthetic rate of cucumbers can be enhanced by a variety of light intensities. The net photosynthetic rate of cucumbers attains its zenith under conditions of CO□ concentrations ranging from 1200-1600 ppm, in conjunction with light intensities spanning from 500-1000 μmol·m^-2^·s^-2^ (Fig. 3). This phenomenon can be attributed to the capacity of elevated CO□ levels to elevate the light saturation point of plants. In the presence of optimal light intensity, higher levels of light energy conversion efficiency are exhibited by plants. Nonetheless, in conditions of low light intensity or intense light conditions, insufficient supplies of ATP and NADPH act as a limiting factor in the enhancement of net assimilation efficiency(Xin *et al*., 2019). An increase in light intensity and carbon dioxide concentration has been shown to progressively enhance growth parameters such as fresh weight, dry weight and leaf area. The interaction between light intensity and carbon dioxide concentration can modify biomass distribution during growth(Hosseinzadeh *et al*., 2021). The findings of the present study demonstrate that carbon dioxide enrichment and high light intensity significantly increased the total biomass and yield of cucumbers. Furthermore, it was observed that carbon dioxide enrichment had a more pronounced effect on root development.

In conditions that are conducive to photosynthesis, an increase in the concentration of carbon dioxide, in conjunction with an escalation in light intensity, has been observed to exert a substantial influence on the growth parameters of plants, including plant height, leaf area, and biomass(Pan *et al*., 2020). The present study demonstrates that the yield of cucumber is significantly increased under eCO_2_ conditions, especially at higher light intensities. This phenomenon is attributed to the efficient flow of carbon between source and sink compartments, facilitated by the addition of carbon dioxide and increased light intensity. The effective photosynthetic radiation in greenhouses is significantly lower than in the external environment. Consequently, the integration of supplemental carbon dioxide with augmented light intensity is poised to emerge as a pivotal strategy for augmenting crop yield and quality(Prakash *et al*., 2023; Kim *et al*., 2025).

### 4.3 The effect of temperature on cucumber photosynthesis, growth and yield

Temperature exerts a regulatory influence on the growth rate and morphogenesis of plants, with this influence being exerted through the medium of the activity of enzymes related to photosynthesis, the photorespiration rate, and the efficiency of carbon allocation(Rachmilevitch *et al*., 2006; Amaral *et al*., 2024). Within the appropriate temperature range, increased carbon dioxide concentrations typically enhance photosynthetic rates and promote biomass accumulation(Sharma *et al*., 2020). However, at excessively high temperatures, photosynthesis may be inhibited, altering the composition of secondary metabolites, which may be difficult to restore even under high carbon dioxide concentrations(Roy *et al*., 2024). Conversely, low temperatures may lead to cellular damage, metabolic slowdown, and growth stagnation(Cao *et al*., 2024). A meta-analysis of cucumbers cultivated under conditions of elevated carbon dioxide concentrations has demonstrated that net photosynthetic rates exhibit a significant increase in response to rising temperatures, thereby promoting biomass and yield accumulation. As a thermophilic plant, cucumbers exhibit increased photosynthetic rates with rising temperatures within a certain temperature range. Additionally, as carbon dioxide concentrations rise, the optimal photosynthetic temperature for crops also increases(Nilsen *et al*., 1983). Overall, both eCO_2_ and elevated temperatures have positive effects on cucumber biomass accumulation and yield. Surprisingly, we found that at carbon dioxide concentrations of 400-800 ppm, higher temperatures (24–32°C) had a less pronounced effect on yield enhancement compared to lower temperatures (Fig. 7). A comparison of the available data sets revealed a positive correlation between yield and both warming and eCO_2_ for the majority of crops, whilst the yield of corn was found to be negatively impacted(Shin *et al*., 2022; Zhu *et al*., 2023). Contrary to expectations, the findings revealed that at carbon dioxide concentrations ranging from 400-800 ppm, elevated temperatures (24–32°C) exerted a comparatively diminished influence on the enhancement of cucumber yield in comparison to reduced temperatures. This may be because the eCO_2_ level is not high enough, so even though photosynthetic rates and total biomass accumulation increase at higher temperatures, accelerated phenological shifts and source-sink imbalances prevent photosynthetic products from being effectively transported to economically productive parts of the plant. However, the overall impact of eCO_2_ and elevated temperature levels on cucumber biomass accumulation and yield is positive, with these factors playing a significant role in the regulation of photosynthesis, growth, and yield. This indicates that they remain a highly reliable regulatory strategy in production.

### 4.4 The effect of humidity on cucumber photosynthesis, growth and yield

The primary function of air humidity is to regulate transpiration rates and the rate at which CO□ enters leaves by affecting stomatal conductance. Moderate increases in humidity have been demonstrated to promote stomatal opening and increase carbon dioxide absorption, thereby improving photosynthetic efficiency(Urban *et al*., 2017). Higher humidity at eCO_2_ increased the net photosynthetic rate of cucumbers, but we found that this was not the most favourable combination for biomass accumulation and yield improvement. eCO_2_ and higher humidity increased the net photosynthetic rate of cucumbers, but we found that this was not the most favourable combination for biomass accumulation and yield improvement. While higher humidity levels have been demonstrated to enhance water status and augment the net photosynthetic rate, they have also been observed to exert deleterious effects. On the one hand, excessive humidity reduced plant transpiration rate, affecting nutrient absorption and transport(McDonald *et al*., 2002), and even causing stress to plants when combined with high temperatures(Cornish *et al*., 2025). the other hand, high humidity may also increase the risk of plant diseases, further affecting plant health and yield(Li *et al*., 2022). Moreover, while carbon dioxide enrichment has been demonstrated to enhance the drought resistance of plants, it has also been observed to prompt accelerated transpiration in cucumbers under conditions of low humidity. This phenomenon occurs as a protective response, wherein stomata close to prevent excessive water loss, consequently restricting carbon dioxide absorption and reducing photosynthetic efficiency(Van De Sanden and Veen, 1992). Consequently, the process of enriching carbon dioxide with the objective of enhancing productivity ought to be executed with meticulous humidity regulation.

### 4.5 The effect of fertilisation on cucumber photosynthesis, growth and yield

Fertilisers play a pivotal role in facilitating the optimal growth and development of vegetables, with particular emphasis on essential nutrients such as nitrogen, phosphorus, and potassium. In high-concentration CO_2_ environments, the nutrient use efficiency of plants undergoes changes(Cui *et al*., 2023). Nitrogen is a pivotal component of chlorophyll and photosynthetic enzymes. An adequate N supply is imperative for sustaining photosynthetic efficiency under conditions of CO□ enrichment, and can substantially augment photosynthetic rates and biomass under such conditions(Hazra *et al*., 2019). It is evident that an increase in carbon dioxide concentrations exerts a substantial influence on the photosynthetic rates, biomass accumulation, and yield of cucumbers, when different nitrogen fertiliser supplies are employed. A substantial increase in the photosynthetic rate, biomass accumulation, and yield of cucumbers was observed in experiments conducted with higher nitrogen supply levels. However, the supply of low nitrogen had a minimal promotional effect on cucumbers under eCO_2_ conditions, and even inhibited biomass accumulation at carbon dioxide concentrations of 400-800 ppm. Furthermore, at concentrations ranging from 400-800 ppm and from 1200-1600 ppm, an inhibitory effect on yield was observed. This is because although carbon dioxide enrichment usually stimulates carbon assimilation, it can also lead to a decrease in nutrient concentrations in plant tissues, especially nitrogen(Zhao *et al*., 2021). In instances where the supply of nitrogen fertiliser is inadequate, the promoting effect of elevated carbon dioxide concentrations on plant growth is counteracted(Reich *et al*., 2014).

Phosphorus is imperative for energy transfer, root development, and flower and fruit formation, while potassium is involved in regulating stomatal opening and closing, enzyme activation, and water relations(Gu *et al*., 2024; Jan *et al*., 2024). Increased carbon dioxide concentrations can reduce the accumulation of phosphorus, an essential element for plants(Bouain *et al*., 2022). In circumstances where phosphorus supply is inadequate, elevated carbon dioxide concentrations have been demonstrated to exert a deleterious effect on photosynthesis, potentially weakening or even preventing it from occurring(Pandey *et al*., 2015). Increased carbon dioxide concentrations can reduce the accumulation of phosphorus, an essential element for plants. When phosphorus fertiliser supply is insufficient, the promotional effect of elevated carbon dioxide concentrations on plant growth may weaken or even inhibit photosynthesis. In summary, under eCO_2_ conditions, the application of an appropriate phosphorus fertiliser concentration in the nutrient solution has been demonstrated to promote cucumber photosynthesis and yield, with higher phosphorus concentrations having a significant effect on biomass accumulation. It is noteworthy that within the range of 800-1200 ppm of carbon dioxide and a phosphorus content in the nutrient solution that corresponds to the standard value (1 mmol·L^-1^), there is a promotion of approximately 50% in the photosynthesis, biomass, and yield of cucumbers. Potassium plays a pivotal role in plants by regulating osmosis, activating enzymes, and controlling stomatal movement. By influencing photosynthesis, it ensures the normal process of carbon assimilation, thereby regulating plant growth and yield(Ho *et al*., 2020). Although higher potassium fertiliser concentrations can significantly increase cucumber photosynthetic rates, their promotional effects on biomass and yield are less pronounced than those of conventional nutrient solution concentrations. This may be because high-potassium nutrient solutions interfere with plants’ absorption and utilisation of other essential elements, leading to nutrient imbalance and consequently reducing plant biomass and yield(Ho *et al*., 2020).

### 4.6 Limitations of the study—future research

The present meta-analysis examined the response of cucumber photosynthesis, growth, and yield under carbon dioxide enrichment. However, the availability of eCO_2_ data was limited, and no detailed gradient interaction analysis was conducted on the effects of eCO_2_ and other environmental factors (light intensity, temperature, etc.) on cucumbers. Furthermore, the paucity of data precluded analysis of the effects of eCO_2_ on cucumber quality, a subject that demands further research in the future. Consequently, further data is required to facilitate precise predictions regarding the impact of eCO_2_ on cucumber growth and yield in the context of global climate change and greenhouse conditions characterised by microclimates.

## 5. Conclusion

The findings of this study suggest that the promotion of photosynthesis, growth, and yield in cucumbers under eCO_2_ conditions is of considerable significance. The findings of this study demonstrate that elevated carbon dioxide concentrations have a positive impact on net photosynthesis, biomass, and yield, with an observed increase of 56.31%, 27.75%, and 21.98%, respectively. The optimal effect was observed within a concentration range of 800-1200 ppm. Furthermore, for eCO_2_, it has been demonstrated that elevated light intensity, temperature, and adequate humidity and fertiliser supply interact synergistically to create optimal conditions for cucumber production. The present study analysed the growth response of cucumbers under carbon dioxide enrichment, which will provide a more informed theoretical framework for environmental regulation in cucumber cultivation and contribute to green, sustainable development.

## Acknowledgments

This research was funded by the Sichuan Provincial Natural Science Foundation Project (2024NSFSC0397); the Agricultural Science and Technology Innovation Program (ASTIP-CAAS, 34-IUA-03); the Local Financial Funds of National Agricultural Science and Technology Center, Chengdu (NASC2022KR01, NASC2023ST06, NASC2024KR03).

## Author contributions

Conceptualization: X.L.; writing—review and editing: X.L.; Q.L.; B.L.; Y.X.; Z.W.; S.L.; and Q.S.; visualization: X.L.; X.L.; funding acquisition: Q.L. All authors have read and agreed to the published version of the manuscript.

## Data availability

The datasets generated and analyzed during the current study are available from the sources cited and, if necessary, from the corresponding author on request.

## References

Abdeldaym EA, Hassan HA, El-Mogy MM, Mohamed MS, Abuarab ME, Omar HS. 2024. Elevated concentrations of soil carbon dioxide with partial root-zone drying enhance drought tolerance and agro-physiological characteristics by regulating the expression of genes related to aquaporin and stress response in cucumber plants. BMC Plant Biology 24, 917.

Amaral J, Lobo AKM, Carmo□Silva E. 2024. Regulation of Rubisco activity in crops. New Phytologist 241, 35–51.

Bouain N, Cho H, Sandhu J, Tuiwong P, Prom-u-thai C, Zheng L, Shahzad Z, Rouached H. 2022. Plant growth stimulation by high CO2 depends on phosphorus homeostasis in chloroplasts. Current Biology 32, 4493-4500.e4.

Burda BU, O’Connor EA, Webber EM, Redmond N, Perdue LA. 2017. Estimating data from figures with a Web□based program: Considerations for a systematic review. Research Synthesis Methods 8, 258–262.

Cao J, Bao J, Lan S, Qin X, Ma S, Li S. 2024. Research progress on low-temperature stress response mechanisms and mitigation strategies in plants. Plant Growth Regulation 104, 1355–1376.

Chen Z, Kang X, Nie H, Zheng S, Zhang T, Zhou D, Xing G, Sun S. 2019. Introduction of Exogenous Glycolate Catabolic Pathway Can Strongly Enhances Photosynthesis and Biomass Yield of Cucumber Grown in a Low-CO2 Environment. Frontiers in Plant Science 10, 702.

Chen Z-F, Wang T-H, Feng C-Y, Guo H-F, Guan X-X, Zhang T-L, Li W-Z, Xing G-M, Sun S, Tan G-F. 2022. Multigene manipulation of photosynthetic carbon metabolism enhances the photosynthetic capacity and biomass yield of cucumber under low-CO2 environment. Frontiers in Plant Science 13, 1005261.

Cornish AE, Kooperman GJ, Grundstein AJ, Skinner CB, Swann ALS. 2025. The impacts of plant physiological responses to rising CO2 on humidity-based extreme heat. npj Climate and Atmospheric Science 8, 167.

Cui E, Xia J, Luo Y. 2023. Nitrogen use strategy drives interspecific differences in plant photosynthetic CO_2_ acclimation. Global Change Biology 29, 3667–3677.

Dong J, Gruda N, Li X, Tang Y, Zhang P, Duan Z. 2020. Sustainable vegetable production under changing climate: The impact of elevated CO2 on yield of vegetables and the interactions with environments-A review. Journal of Cleaner Production 253, 119920.

Dong J, Li X, Nazim G, Duan Z. 2018. Interactive effects of elevated carbon dioxide and nitrogen availability on fruit quality of cucumber (Cucumis sativus L.). Journal of Integrative Agriculture 17, 2438–2446.

Du Y, Liu X, Zhang L, Zhou W. 2023. Drip irrigation in agricultural saline-alkali land controls soil salinity and improves crop yield: Evidence from a global meta-analysis. Science of The Total Environment 880, 163226.

Gu H, Li J, Lu Z, Li X, Cong R, Ren T, Lu J. 2024. Effects of combined application of nitrogen and potassium on oil concentration and fatty acid component of oilseed rape (Brassica napus L.). Field Crops Research 306, 109229.

Hao P-F, Qiu C-W, Ding G, Vincze E, Zhang G, Zhang Y, Wu F. 2020. Agriculture organic wastes fermentation CO2 enrichment in greenhouse and the fermentation residues improve growth, yield and fruit quality in tomato. Journal of Cleaner Production 275, 123885.

Hazra S, Swain DK, Bhadoria PBS. 2019. Wheat grown under elevated CO2 was more responsive to nitrogen fertilizer in Eastern India. European Journal of Agronomy 105, 1–12.

Hedges LV, Gurevitch J, Curtis PS. 1999. The meta-analysis of response ratios in experimental ecology. Ecology 80, 1150–1156.

Ho L-H, Rode R, Siegel M, Reinhardt F, Neuhaus HE, Yvin J-C, Pluchon S, Hosseini SA, Pommerrenig B. 2020. Potassium Application Boosts Photosynthesis and Sorbitol Biosynthesis and Accelerates Cold Acclimation of Common Plantain (Plantago major L.). Plants 9, 1259.

Horton P, Long SP, Smith P, Banwart SA, Beerling DJ. 2021. Technologies to deliver food and climate security through agriculture. Nature Plants 7, 250–255.

Hosseinzadeh M, Aliniaeifard S, Shomali A, Didaran. 2021. Interaction of Light Intensity and CO2 Concentration Alters Biomass Partitioning in Chrysanthemum. Journal of Horticultural Research 29, 45–56.

Hou L, Shang M, Chen Y, Zhang J, Xu X, Song H, Zheng S, Li M, Xing G. 2021. Physiological and molecular mechanisms of elevated CO2 in promoting the growth of pak choi (Brassica rapa ssp. chinensis). Scientia Horticulturae 288, 110318.

Huang B, Xu Y. 2015. Cellular and Molecular Mechanisms for Elevated CO_2_ –Regulation of Plant Growth and Stress Adaptation. Crop Science 55, 1405–1424.

IPCC. 2023. Sections. In: Climate Change (2023). Synthesis Report. Contribution of working groups I, II and III to the Sixth Assessment Report of the Intergovernmental Panel on Climate Change [Lee H, Romero J (eds.)]. Geneva, Switzerland, 35–115.

Jan AL, Amanullah, Mihoub A, Fawad M, Saeed MF, Khan I, Radicetti E, Jamal A. 2024. Enhancing wheat performance through phosphorus and zinc management strategies under varied irrigation regimes. Environment, Development and Sustainability

Kim D, Goo H, Yoon S, Yoon J, Kim H, Yoo Y, Lee M, Park KS. 2025. Effects of Supplemental LED Lighting based on Solar Irradiation and Carbon Dioxide Enrichment on Photosynthesis, Growth, and Yield of Greenhouse-Grown Tomato Plants. 28 1, 1–12.

Kläring H-P, Hauschild C, Heißner A, Bar-Yosef B. 2007. Model-based control of CO2 concentration in greenhouses at ambient levels increases cucumber yield. Agricultural and Forest Meteorology 143, 208–216.

Lajeunesse MJ. 2016. Facilitating systematic reviews, data extraction and meta□analysis with the METAGEAR package for R. (R Fitzjohn, Ed.). Methods in Ecology and Evolution 7, 323–330.

Li T, Zhou J, Liu R, Yuan Z, Li J. 2022. Effects of photo-selective nets and air humidity coupling on tomato resistance to Botrytis cinerea. Scientia Horticulturae 305, 111356.

Li Y, Zhu S, Hu J, Guo S, Shi H, Cao Y. 2025. Effects of Different Densities of Carbon Dioxide Generation Bags on Cucumber Growth and Yield. Horticulturae 11, 218.

Liu Y, Sun S, Xing G, Li J, Zhang Z, Yuan H, Zheng J. 2018. Effects of different concentrations of CO_2_ on the growth and yield of greenhouse cucumber Cucumis sativus L. Journal of Shanxi Agricultural University (Natural Science Edition) 38, 53–58.

Long SP, Ainsworth EA, Leakey ADB, Nösberger J, Ort DR. 2006. Food for Thought: Lower-Than-Expected Crop Yield Stimulation with Rising CO_2_ Concentrations. Science 312, 1918–1921.

Maschler J, Bialic□Murphy L, Wan J, et al. 2022. Links across ecological scales: Plant biomass responses to elevated CO_2_. Global Change Biology 28, 6115–6134.

McDonald EP, Erickson JE, Kruger EL. 2002. Research note: Can decreased transpiration limit plant nitrogen acquisition in elevated CO2? Functional Plant Biology 29, 1115.

Nilsen S, Hovland K, Dons C, Sletten SP. 1983. Effect of CO2 enrichment on photosynthesis, growth and yield of tomato. 20, 1–14.

Osman M, Qaryouti M, Alharbi S, Alghamdi B, Al-Soqeer A, Alharbi A, Almutairi K, Abdelaziz ME. 2024. Impact of CO2 Enrichment on Growth, Yield and Fruit Quality of F1 Hybrid Strawberry Grown under Controlled Greenhouse Condition. Horticulturae 10, 941.

Pan T, Wang Y, Wang L, Ding J, Cao Y, Qin G, Yan L, Xi L, Zhang J, Zou Z. 2020. Increased CO_2_ and light intensity regulate growth and leaf gas exchange in tomato. Physiologia Plantarum 168, 694–708.

Pandey R, Zinta G, AbdElgawad H, Ahmad A, Jain V, Janssens IA. 2015. Physiological and molecular alterations in plants exposed to high [CO2] under phosphorus stress. Biotechnology Advances 33, 303–316.

Prakash V, Lunagaria MM, Trivedi AP, Upadhyaya A, Kumar R, Das A, Kumar Gupta A, Kumar Y. 2023. Shading and PAR under different density agrivoltaic systems, their simulation and effect on wheat productivity. European Journal of Agronomy 149, 126922.

Qiang Q, Gao Y, Yu B, Wang M, Ni W, Li S, Zhang T, Li W, Lin L. 2020. Elevated CO2 enhances growth and differentially affects saponin content in Paris polyphylla var. yunnanensis. Industrial Crops and Products 147, 112124.

Rachmilevitch S, Huang B, Lambers H. 2006. Assimilation and allocation of carbon and nitrogen of thermal and nonthermal Agrostis species in response to high soil temperature. New Phytologist 170, 479–490.

Reich PB, Hobbie SE, Lee TD. 2014. Plant growth enhancement by elevated CO2 eliminated by joint water and nitrogen limitation. Nature Geoscience 7, 920–924.

Robredo A, Pérez-López U, De La Maza HS, González-Moro B, Lacuesta M, Mena-Petite A, Muñoz-Rueda A. 2007. Elevated CO2 alleviates the impact of drought on barley improving water status by lowering stomatal conductance and delaying its effects on photosynthesis. Environmental and Experimental Botany 59, 252–263.

Roy S, Kapoor R, Mathur P. 2024. Revisiting Changes in Growth, Physiology and Stress Responses of Plants under the Effect of Enhanced CO2 and Temperature. Plant And Cell Physiology 65, 1–3.

Sharma S, Walia S, Rathore S, Kumar P, Kumar R. 2020. Combined effect of elevated CO2 and temperature on growth, biomass and secondary metabolite of Hypericum perforatum L. in a western Himalayan region. Journal of Applied Research on Medicinal and Aromatic Plants 16, 100239.

Shin J, Hwang I, Kim D, Kim J, Kim JH, Son JE. 2022. Waning advantages of CO2 enrichment on photosynthesis and productivity due to accelerated phase transition and source-sink imbalance in sweet pepper. Scientia Horticulturae 301, 111130.

Sugiura D, Wang Y, Kono M, Mizokami Y. 2024. Exploring the responses of crop photosynthesis to CO2 elevation at the molecular, physiological, and morphological levels toward increasing crop production. Crop and Environment 3, 75–83.

Syed AM, Hachem C. 2019. Review of Design Trends in Lighting, Environmental Controls, Carbon Dioxide Supplementation, Passive Design, and Renewable Energy Systems for Agricultural Greenhouses. Journal of Biosystems Engineering 44, 28–36.

Urban J, Ingwers MW, McGuire MA, Teskey RO. 2017. Increase in leaf temperature opens stomata and decouples net photosynthesis from stomatal conductance in Pinus taeda and Populus deltoides x nigra. Journal of Experimental Botany 68, 1757–1767.

Van De Sanden PACM, Veen BW. 1992. Effects of air humidity and nutrient solution concentration on growth, water potential and stomatal conductance of cucumber seedlings. Scientia Horticulturae 50, 173–186.

Wang L, Feng Z, Schjoerring JK. 2013. Effects of elevated atmospheric CO2 on physiology and yield of wheat (Triticum aestivum L.): A meta-analytic test of current hypotheses. Agriculture, Ecosystems & Environment 178, 57–63.

Wang X, Wang L, Chen Y, Hu Y, Guan R, Li M, Wang L, Zhang Y. 2024. Mitigating the negative effect of warming on crop yield: Assessing the carbon fertilization and organic amendment application effect. Field Crops Research 311, 109370.

Xin P, Li B, Zhang H, Hu J. 2019. Optimization and control of the light environment for greenhouse crop production. Scientific Reports 9, 8650.

Zhang Z, Liu L, Zhang M, Zhang Y, Wang Q. 2014. Effect of carbon dioxide enrichment on health-promoting compounds and organoleptic properties of tomato fruits grown in greenhouse. Food Chemistry 153, 157–163.

Zhang Y, Yasutake D, Hidaka K, Kitano M, Okayasu T. 2020. CFD analysis for evaluating and optimizing spatial distribution of CO2 concentration in a strawberry greenhouse under different CO2 enrichment methods. Computers and Electronics in Agriculture 179, 105811.

Zhao H-L, Chang T-G, Xiao Y, Zhu X-G. 2021. Potential metabolic mechanisms for inhibited chloroplast nitrogen assimilation under high CO2. Plant Physiology 187, 1812–1833.

Zhao H, Zhai X, Guo L, Yang Y, Li J, Ren C, Wang K, Liu X, Zhan R, Wang K. 2019. Comparing protected cucumber and field cucumber production systems in China based on emergy analysis. Journal of Cleaner Production 236, 117648.

Zhu C, Wolf J, Zhang J, Anderegg WRL, Bunce JA, Ziska LH. 2023. Rising temperatures can negate CO2 fertilization effects on global staple crop yields: A meta-regression analysis. Agricultural and Forest Meteorology 342, 109737.

Ziska LH, Bunce JA. 2007. Predicting the impact of changing CO_2_ on crop yields: some thoughts on food. New Phytologist 175, 607–618.

